# Engrafted Human Induced Pluripotent Stem Cell-Derived Cardiomyocytes Undergo Clonal Expansion *In Vivo*

**DOI:** 10.1101/847665

**Authors:** Danny El-Nachef, Darrian Bugg, Kevin M. Beussman, Amy M. Martinson, Charles E. Murry, Nathan J. Sniadecki, Jennifer Davis

**Author notes:** **Correspondence** Jennifer Davis. Mail: 850 Republican Street, Building D, Room 343, Seattle, WA 98109.

## Abstract

Preclinical studies have suggested that transplanted human pluripotent stem cell-derived cardiomyocyte (hPSC-CM) grafts expand due to proliferation. This knowledge came from cell cycle activity measurements that cannot discriminate between cytokinesis or DNA synthesis associated with hypertrophy. To refine our understanding of hPSC-CM cell therapy, we genetically engineered a cardiomyocyte-specific fluorescent barcoding system into an hPSC line. Since cellular progeny have the same color as parental hPSC-CMs, we could identify subsets of engrafted hPSC-CMs that clonally expanded, with the remainder being non-proliferative and hypertrophic.

## Main Text

Preclinical studies have suggested that transplanted human pluripotent stem cell-derived cardiomyocyte (hPSC-CM) grafts expand due to proliferation.^1^ This knowledge came from cell cycle activity measurements that cannot discriminate between cytokinesis or DNA synthesis associated with hypertrophy. To refine our understanding of hPSC-CM cell therapy, we genetically engineered a cardiomyocyte-specific fluorescent barcoding system into an hPSC line. Since cellular progeny have the same color as parental hPSC-CMs, we could identify subsets of engrafted hPSC-CMs that clonally expanded, with the remainder being non-proliferative and hypertrophic.

We generated our hPSC line by knocking four copies of the Cre-dependent Brainbow 3.2 lineage reporter^2^ into WTC11 cells (Figure A). These rainbow hPSCs were transduced with cardiac troponin T (cTnT)-driven Cre, which restricts expression of the rainbow barcoding system to committed cardiomyocytes (Figure A). Rainbow-labeling was observed after 7 days of differentiation and immunostaining confirmed the labeled cells expressed cTnT (Figure B). We aimed for sparse labeling so we could track individual cells over time since neighboring cells would not have the same color (Figure C). Cre-mediated recombination elicited all eighteen of the possible hues, but monocolor-labeling dominated (Figure D).

**Figure.**
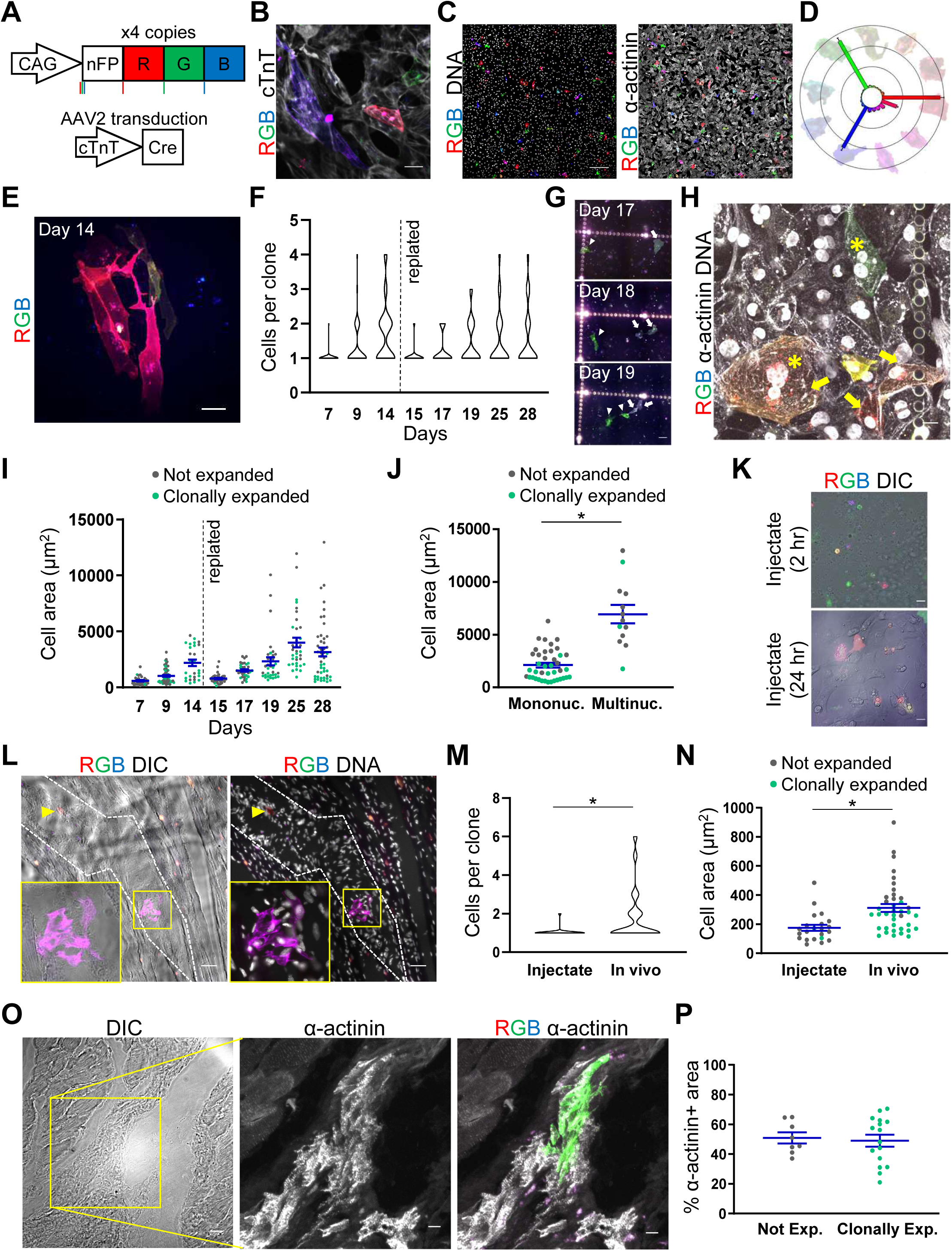
Select cardiomyocytes clonally expand *in vitro* and *in vivo*. **(A)** Schematic of rainbow hPSC line and cTnT-activated labeling. We replaced the neuron-specific Thy1 promoter in the Brainbow 3.2 transgene construct with a universal CAG promoter. This construct was placed between AAVS1 safe harbor locus homology arms and co-electroporated into WTC11 cells with a construct expressing Cas9 and a guide RNA directed to the AAVS1 locus. We selected a WTC11 colony that had four copies of the Brainbow construct – two copies from AAVS1-targeted bi-allelic knock-in and two copies from random insertion. Before expression of Cre recombinase, rainbow hPSCs express a non-fluorescent GFP mutant (nFP). By means of the Cre-lox recombination system, gene excision may occur randomly at either the loxP (red vertical bars), lox2 (green vertical bars), or loxN (blue vertical bars) sites, resulting in expression of one of three fluorescent proteins: mOrange2 (herein referred to as R and colored red), eGFP (herein referred to as G and colored green), and mKate2 (herein referred to as B and colored blue). To label cells after they committed to the cardiomyocyte lineage, we utilized an adeno-associated virus (AAV) plasmid (Addgene 69916) that contained AAV ITR sequences flanking a cardiac troponin T promoter upstream of the *Cre* and *TdTomato* genes. Because TdTomato has spectral overlap with the rainbow reporter, it was excised by NotI digestion and the resulting construct was used for AAV serotype 2 (AAV2) production using standard protocols. AAV2 was added to rainbow hPSCs on day 0 of the cardiomyocyte differentiation. We used a previously established directed differentiation protocol with slight modification.^1^ Briefly, hPSCs were seeded on Matrigel coated vessels (350,000 per well of 6-well plate or 100,000 per well of 24-well plate) in mTeSR media supplemented with 10 µM Y-27632. The next day (day −1), the media was changed to mTeSR media supplemented with 1 µM Chiron 99021 (Cayman Chemicals). On day 0, media was changed to RPMI (Gibco) supplemented with B27 supplement minus insulin (Thermo Fisher) and 5 µM Chiron 99021. On day 2, media was changed to RPMI supplemented with B27 supplement minus insulin and 2 µM Wnt inhibitor Wnt-C59 (Selleckchem). On day 4, media was changed to RPMI supplemented with B27 minus insulin. On day 6, media was changed to RPMI supplemented with B27 supplement plus insulin (cardio media), with recurring media changes every 2-3 days until endpoint. **(B)** Rainbow-labeled hPSC-CMs express cTnT protein, scale bar is 20 µm. On day 14, cells were fixed with 4% PFA, permeabilized/blocked in 5% goat serum in PBS supplemented with 0.1% TritonX-100 and stained for cTnT (ThermoFisher MA5-12960). Imaging was performed on a Nikon TiE equipped with Yokogawa W1 spinning disk confocal system and high sensitivity EMCCD camera with the following settings (excitation fixed-LED laser wavelength (nm), emission filter): eGFP (488, GFP), mOrange2 (561, TRITC), mKate2 (561, Cy5), and cTnT was visualized with an Alexa Fluor 405 secondary antibody (405 nm laser, DAPI filter). **(C)** Representative image of sparsely-labeled rainbow hPSC-CMs on day 9 and stained for DNA (Hoechst, Life Technologies H3570) and α-actinin (Sigma A7811), scale bar is 200 µm. AAV2 was serially diluted to determine optimal dose for sparse labeling. Images were collected on a Leica SP8 confocal microscope with the following laser and spectral detector settings (excitation laser wavelength (nm), emission spectral detector range (nm): eGFP (488, 498-522), mOrange2 (552, 563-590), mKate2 (552, 633-638), Hoechst (405, 410-480), and α-actinin was detected with an Alexa Fluor 647 secondary antibody (638, 657-720). **(D)** Resultant images were analyzed on a per pixel basis and binned by the eighteen possible hues using a custom generated MATLAB code. Examples of rainbow hPSC-CMs expressing different hues are shown around the color wheel. The number of unique hues was determined as follows: copy number analysis using qPCR of rainbow hPSC genomic DNA demonstrated a total of four copies of the Brainbow cassette. Because there are three possible fluorescent proteins that can be expressed from each cassette, the number of unique possible outcomes can be described by the trinomial theorem, [(*n*+2)(*n*+1)]/2, where *n* = the number of copies of the cassette. To account for the possibility that a cell can undergo 1, 2, 3, or 4 recombination events this equation was solved for n=1, n=2, n=3, and n=4, and summed, indicating 34 unique recombination outcomes. However, twelve of the unique combinations result in duplicated colors that vary only by intensity (i.e. one copy of red versus four copies of red result in the same hue, though with a different signal intensity) and another four unique outcomes result in duplicated hues that vary only by tint (i.e. one copy of red versus one copy of green and one copy of blue with two copies of red results in the same red hue, though with a different tint of white). Thus, there are eighteen unique hues that can be represented on a color wheel. The primary colors were most represented, presumably due to the likelihood of only one Brainbow copy undergoing Cre-lox recombination and the greater proportion of recombination outcomes resulting in primary color expression. **(E)** Example of clonally expanded rainbow hPSC-CMs at day 14 in the differentiation, scale bar is 10 µm. **(F)** Quantification of rainbow hPSC-CM clonal expansion over time and after replating on day 14. **(G)** Time lapse imaging of replated rainbow hPSC-CMs showing cell proliferation, scale bar is 40 µm. Arrow points to a hPSC-CM that divides between day 17 and 18, arrowhead points to a hPSC-CM that divides between day 18 and 19. Single cell suspensions were generated by incubating the culture with a 1:1 mixture of 0.25% Trypsin and Versene for 7 minutes at 37° C. Digestion was stopped with Stop Solution (cardio media with 10% FBS and 0.2 Kunitz units/µL DNase I) and cells were replated on a coordinate gridded well (Ibidi) in cardio media supplemented with 10 µM Y-27632 and 5% FBS for the first 24 hours, then returned to standard cardio media and replaced every 2-3 days. **(H)** Staining for α-actinin and nuclei of rainbow hPSC-CMs on day 28 shows instances of clonal expansion and multinucleation, scale bar is 20 µm. Arrows point to clonally expanded hPSC-CMs, asterisks indicate multinucleated hPSC-CMs. **(I)** Non-expanded (i.e. singlets) and clonally expanded hPSC-CMs (i.e. clones of two or more cells) are color coded gray or green, respectively. Cells were manually traced and counted in ImageJ. Cell area was determined using the ‘measure’ feature in ImageJ. **(J)** Quantification of cell area as a function of the number of nuclei per cell for rainbow hPSC-CMs on day 28. Non-dividing and clonally expanded hPSC-CMs are labeled in gray or green, respectively, * indicates p < 0.05. **(K)** Images of the injectate after 2 and 24 hours after preparation show that there is sparse labeling of rainbow hPSC-CMs, scale bar is 20 µm. On day 14, sparsely-labeled rainbow hPSC-CMs were resuspended in Matrigel supplemented with a prosurvival cocktail consisting of 10 μM ZVAD-FMK/caspase inhibitor (EMD-Millipore); 50 nM TAT-BH4/BCL-XL (EMD-Millipore); 200 nM cyclosporine A (Novartis); 50 μM pinacidil (Sigma); 10 μM Y-27632, and 100 ng/ml IGF-1 (Peprotech) at a concentration of 50 million cells/mL. **(L)** Imaging of host rat heart with engrafted hPSC-CMs at 14 days after the procedure, graft region is marked by dashed, white lines. Arrowhead points to a singlet rainbow hPSC-CM, box and inset show clonally expanded hPSC-CM, scale bar is 40 µm. All animal experiments were performed in accordance with the Guide for the Care and Use of Laboratory Animals published by the US National Institute of Health and was approved by the University of Washington institutional animal care and use committee (IACUC protocol # 4376-01). Athymic male Sprague Dawley rats (rnu-rnu, 250–300 g, Harlan) were immunosuppressed using cyclosporin at a dose of 5 mg/kg for a duration of seven days starting a day prior to the engraftment procedure. On the day of the engraftment procedure, animals received 1 mg/kg sustained-release buprenorphine and were induced with inhaled isoflurane and mechanically ventilated. A left lateral thoracotomy was performed to expose the left ventricle where three injections of equal size from the base to apex were preformed to deliver 5 million cells in total. Fourteen days after the engraftment procedure, the rats were euthanized with Euthasol, their hearts were collected in KB buffer (20 mM KCl, 10 mM KH_2_PO_4_, 70 mM K^+^-glutamate, 1 mM MgCl_2_, 25 mM glucose, 20 mM taurine, 0.5 mM EGTA, 10 mM HEPES, 0.1% albumin, pH 7.4 with KOH), cannulated and perfused with KB followed by 4% PFA, additionally fixed with 4% PFA overnight, washed with PBS, mounted in OCT (Tissue-Tek) for cryosectioning of 10 µm thick specimens that were subsequently stained and analyzed using the methodology and reagents described above. Quantification **(M)** clonal expansion and **(N)** cell area in engrafted rainbow hPSC-CMs at 14 days after engraftment and comparison to injectate. * indicates p < 0.05. **(O)** Example of α-actinin expression in engrafted rainbow hPSC-CM that clonally expanded *in vivo*, scale bar is 20µm, inset bar is 10 µm. **(P)** Quantification of α-actinin content normalized by cell area for engrafted rainbow hPSC-CMs that did not divide or clonally expanded. Images were analyzed in ImageJ by converting to a binary image using the ‘threshold’ function to quantify the area of α-actinin and then dividing by the total cell area, which was manually traced using the ‘measure’ function. GraphPad Prism V8 was used for two-tailed unpaired t-tests (Figures J, M, N, and P) and one-Way ANOVA with Tukey posthoc tests (Figures F and I). A p-value lower than 0.05 was considered statistically significant. Each quantification was derived from a minimum of three independent experiments (different passages for *in vitro* experiments or different animals in *in vivo* studies). All average values described in the text and figures are mean values and error bars represent standard error of the mean.

By day 14, we observed that hPSC-CMs had clonally expanded (Figure E). On average, the number of cardiomyocytes per clone went from 1.03 to 1.71 (day 7 versus 14, p<0.001, Figure F). While most rainbow hPSC-CMs did not proliferate over time, some were highly proliferative and a subset of hPSC-CMs continued to proliferate after replating at day 14 (Figure F). Repeat imaging of the replated hPSC-CMs over days 15-28 confirmed that (1) neighboring cells had unique hues (Figures F and G) and (2) all daughter cardiomyocytes inherited the parental fluorescent-barcode (Figure G), definitively demonstrating these clusters arise from clonal expansion (Figure H). We noted that hPSC-CM underwent hypertrophic growth as measured by cell area (Figure I), and by day 28, clonally expanded hPSC-CMs were 4.07-fold smaller than non-dividing hPSC-CMs (p<0.0001, Figure I). Staining reconfirmed cTnT-driven labeling was specifically induced in cardiomyocytes and showed multinucleation in non-dividing, hypertrophied hPSC-CMs (Figures H and J).

For the transplantation studies, sparsely-labeled day 14 hPSC-CMs were dissociated, resuspended in Matrigel with prosurvival cocktail, and transplanted into the hearts of immune-compromised athymic rats.^1^ Hearts were harvested two weeks after hPSC-CM engraftment, which equates to the 28-day timepoint *in vitro*. Most engrafted hPSC-CMs hypertrophied (1.75-fold increase versus injectate, p<0.002) while some subsets clonally expanded with the average number of cells per clone increasing to 1.77 (p<0.03, Figures K-O). Twenty-two clusters of engrafted hPSC-CM clones were identified, of those 36.4% had undergone at least one cell division and 18.2% had divided multiple times. To ensure that the results were not driven by false positives, the cell injectate was imaged to confirm that neighboring hPSC-CMs did not express the same barcode at the time of transplantation (Figures K and M). Notably, the heterogeneous proliferative potential among engrafted hPSC-CMs, which would not have been observed without the rainbow single-cell reporter, was not due to differences in sarcomere content, as the proportion of α-actinin+ area was the same in both clonally expanding and non-expanding groups (p>0.77, Figures O-P).

As hPSC-CM therapy is rapidly approaching clinical use, it is critical to understand how these cells behave *in vivo*. Indeed single cell transcriptomics assays^3^ and DNA content analysis^4^ have revealed profound molecular heterogeneity among these hPSC-CMs. It remains unclear how these variances translate to functional heterogeneity, but by generating a cTnT lineage rainbow reporter we could longitudinally track the hypertrophic and proliferative growth of individual hPSC-CMs. This approach demonstrated for the first time that hPSC-CMs have heterogeneous levels of proliferation *in vitro* and after engraftment in host myocardium, revealing a dichotomy between non-dividing and clonally expanding hPSC-CMs in their capacity to hypertrophy. By examining the generation of newly formed cardiomyocytes, rather than utilizing proxies for cell proliferation, this study distinguished *bona fide* cardiomyocyte division versus incomplete cell cycle activation, which has been a source of controversy in the field. Moreover, this analysis found instances of multinucleation that could have led to false identification of cell proliferation with gold standard DNA synthesis or mitosis markers that measure newly formed genomes or nuclei rather than cell division. While the heterogenous proliferative capacity among hPSC-CMs was initially surprising to us, our results are consistent with findings that demonstrated only a few clonally dominant cardiomyocytes give rise to most of the adult zebrafish heart^5^, suggesting a similar mechanism may underlie hPSC-CM graft expansion. The finding that proliferative hPSC-CM express normal levels of sarcomere contractile elements suggests increasing hPSC-CM clonal expansion will more efficiently repopulate myocardium lost to injury. This is in line with cardiac cell therapy optimizations, such as co-transplantation of hPSC-CMs with epicardial cells^1^, demonstrating significantly improved outcomes from stimulating grafted hPSC-CM proliferation. Thus, controlling engrafted hPSC-CM clonal expansion holds great promise for improving cardiac regenerative therapies.

## Article Information

Data sharing: All data and materials will be available from the corresponding author by request.

## References

1. Bargehr J, Ong LP, Colzani M, Davaapil H, Hofsteen P, Bhandari S, Gambardella L, Le Novère N, Iyer D, Sampaziotis F, Weinberger F, Bertero A, Leonard A, Bernard WG, Martinson A, Figg N, Regnier M, Bennett MR, Murry CE, Sinha S. Epicardial cells derived from human embryonic stem cells augment cardiomyocyte-driven heart regeneration. Nat Biotechnol. 2019;37:895–906. DOI: 10.1038/s41587-019-0197-9.

2. Cai D, Cohen KB, Luo T, Lichtman JW, Sanes JR. Improved tools for the Brainbow toolbox. Nat Methods. 2013;10:540–7. DOI: 10.1038/nmeth.2450.

3. Friedman CE, Nguyen Q, Lukowski SW, Helfer A, Chiu HS, Miklas J, Levy S, Suo S, Han J-DJ, Osteil P, Peng G, Jing N, Baillie GJ, Senabouth A, Christ AN, Bruxner TJ, Murry CE, Wong ES, Ding J, Wang Y, Hudson J, Ruohola-Baker H, Bar-Joseph Z, Tam PPL, Powell JE, Palpant NJ. Single-Cell Transcriptomic Analysis of Cardiac Differentiation from Human PSCs Reveals HOPX-Dependent Cardiomyocyte Maturation. Cell Stem Cell. 2018;23:586–598. DOI: 10.1016/j.stem.2018.09.009.

4. Lundy SD, Zhu W-Z, Regnier M, Laflamme MA. Structural and Functional Maturation of Cardiomyocytes Derived from Human Pluripotent Stem Cells. Stem Cells Dev. 2013;22:1991–2002. DOI: 10.1089/scd.2012.0490.

5. Gupta V, Poss KD. Clonally dominant cardiomyocytes direct heart morphogenesis. Nature. 2012;484:479. DOI: 10.1038/nature11045.

